# Incipient parallel evolution of SARS-CoV-2 Deltacron variant in South Brazil

**DOI:** 10.1101/2022.10.06.511203

**Authors:** Fernando Hayashi Sant’Anna, Tiago Finger Andreis, Richard Steiner Salvato, Ana Paula Muterle, Juliana Comerlato, Tatiana Schaffer Gregianini, Regina Bones Barcellos, Fernanda Marques Godinho, Paola Cristina Resende, Gabriel da Luz Wallau, Thaís Regina y Castro, Bruna Campestrini Casarin, Andressa de Almeida Vieira, Alexandre Vargas Schwarzbold, Priscila de Arruda Trindade, Gabriela Luchiari Tumioto Giannini, Luana Freese, Giovana Bristot, Carolina Serpa Brasil, Bruna de Oliveira Rocha, Paloma Bortolini Martins, Francine Hehn de Oliveira, Cock van Oosterhout, Eliana Wendland

**Author notes:** Both authors contributed equally to this manuscript (Fernando Hayashi Sant’Anna, Tiago Finger Andreis). **Corresponding authors:** Eliana Wendland and Cock van Oosterhout, **Address for correspondence:** and.

## Abstract

With the coexistence of multiple lineages and increased international travel, recombination and gene flow are likely to become increasingly important in the adaptive evolution of SARS-CoV-2. This could result in the incipient parallel evolution of multiple recombinant lineages. However, identifying recombinant lineages is challenging, and the true extent of recombinant evolution in SARS-CoV-2 may be underestimated. This study describes the first SARS-CoV-2 Deltacron recombinant case identified in Brazil. We demonstrate that the recombination breakpoint is at the beginning of Spike gene (*S*). The 5′ genome portion (circa 22 kb) resembles the AY.101 lineage (VOC Delta), and the 3′ genome portion (circa 8 kb nucleotides) is most similar to the BA.1.1 lineage (VOC Omicron). Furthermore, evolutionary genomic analyses indicate that the new strain emerged after a single recombination event between lineages of diverse geographical locations in December 2021 in South Brazil. This Deltacron, named AYBA-RS, is one out of almost 30 recombinants described this year. The submission of only four sequences in the GISAID database suggests that this Brazilian lineage had a minor epidemiological impact. On the other hand, the recent emergence of this and various other Deltacron recombinant lineages (i.e., XD, XF, and XS) suggests that gene flow and recombination may play an increasingly important role in the COVID-19 pandemic. We explain the evolutionary and population genetic theory that support this assertion, and we conclude that this stresses the need for continued genomic and epidemiological surveillance. This is particularly important for countries where multiple variants are present, as well as for countries that receive significant inbound international travel.

## Introduction

Similar to other organisms, coronaviruses are subject to three evolutionary forces that can generate genetic variation: mutation, recombination and gene flow. Random mutational changes at the nucleotide level (substitutions, insertions, and deletions) may lead to the emergence of new lineages with increased transmission and immune evasion capacity [1]. During the first two years of the COVID-19 outbreak, most genetic variation was generated by such random mutations, some of which improved the fitness of the virus and enabled it to adapt itself better to its new host and epidemiological environment. Recombination is another source of viral variability that could drive adaptive evolution. In SARS-CoV-2, recombination can occur when distinct variants co-infect the same host cell and exchange genetic material [2,3]. This process is called genetic introgression and it plays an important role in virulence evolution of parasites and pathogens [4]. Gene flow happens when a genotype of a given variant is moved from one population into another. If such gene flow also results in a co-infection in a host already infected with another variant, it might lead to genetic introgression. This is known to have resulted in the evolution of novel subspecies in other human parasites such as *Cryptosporidium spp*. (e.g., [5]), and such subspecies might continue to exchange genetic information that underpins their virulence evolution [6]. In the case of SARS-CoV-2, international travel is likely to contribute considerably to the gene flow of different variants across the globe, thereby increasing the probability of genetic introgression. The combined evolutionary forces of recombination and gene flow are particularly important drivers in pathogen evolution because it enables diverged variants of different geographic locations to exchange genetic variation [4].

The coexistence of multiple SARS-CoV-2 variants can be explained by at least four population genetic or coevolutionary hypotheses. Firstly, multiple variants can be maintained by spatiotemporal variation in selection pressures. Such balancing selection arises when different variants are locally adapted to the epidemiological environment of the geographic region. In that case, no single variant is able to replace all other variants, resulting in the coexistence of multiple variants. This coevolutionary model is known as Red Queen Dynamics [7]. Secondly, SARS-CoV-2 may not yet have reached a coevolutionary equilibrium with its primary host (humans), and novel variants can continue to evolve that can replace the dominant ancestral variant. This would result in the erratic turnover of variants and a spatiotemporal dynamic polymorphism. Thirdly, there is a dynamic coevolutionary equilibrium, with SARS-CoV-2 continuously evolving counter-adaptations to infectious disease control measures (e.g., vaccines) and naturally acquired resistance or tolerance to the disease. The latter is known as a Red Queen Arms Race [7], and it is predicted to lead to a steady turnover of variants. Finally, in a spatially substructured host population, or with multiple reservoir hosts, and with a very large viral population, coexistence of multiple variants is likely, particularly if the fitness of the variants (i.e., their reproductive number, R0) isn’t markedly different. Under each of these scenarios, the global metapopulation of SARS-CoV-2 will possess genetic polymorphism in the form of multiple coexisting variants that are, to some extent, separated in time and/or space. In turn, the presence of such polymorphism implies that gene flow and recombination can rapidly generate genetic novelty, and this can have important evolutionary ramifications [4].

SARS-CoV-2 recombinant lineages can potentially make evolutionary leaps that bridge valleys in the fitness landscape. According to Wright’s Shifting Balance theory (Wright 1982), fitness valleys separate different peaks in the adaptive landscape. The evolutionary path from one adaptive peak to a second even higher peak takes the genotype through a fitness valley. If these intermediate maladapted genotypes have to evolve through a series of single mutations, this may prevent adaptive evolution from ever reaching this higher peak, because the intermediate genotypes are less fit than the ancestor. However, with genetic introgression, multiple substitutions are made in the genomic background all at once, potentially crossing the fitness peak in a single recombination event. In addition, in contrast to random mutation, recombination inserts nucleotide substitutions that have already been “tried and tested” in another genomic background [4].

Adaptations driven by the combined forces of gene flow and recombination could result in the incipient, parallel evolution of multiple recombinant lineages that compete with one another for global dominance. In this hypothetical scenario, recombination offers three potential advantages over mutation: 1) recombination can insert multiple substitutions all at once; 2) these substitutions have been previously selected and are functional in the genomic background of the parental lineage; 3) this enables the recombinant genotype to bridge the fitness valleys in the adaptive fitness landscape and find higher fitness peaks. In addition, increased international travel enables gene flow and recombinant exchange between distinct lineages that have evolved all across the world. Consequently, recombination and gene flow may play an increasingly important role in the transmissibility, severity, and resistance to vaccines and treatments of SARS-CoV-2, and in the evolutionary epidemiology of the COVID-19 pandemic. Here, we examine the evidence for incipient parallel evolution of recombinant lineages, studying SARS-CoV-2 genomes in Brazil. This country has seen less intensive genomic and epidemiological surveillance than other parts of the world, and hence, by studying the SARS-CoV-2 genome sequence variation in Brazil, we may get a better understanding about the extent of cryptic recombinant lineages.

Until September 2022, WHO reported five SARS-CoV-2 variants of concern (VOC): Alpha, Beta, Gamma, Delta, Omicron [8]. The variant Delta (B.1.617.2) emerged in India at the end of 2020 and spread to at least 185 nations [9,10]. WHO then classified this variant as a VOC, given its high transmissibility and potential to cause severe COVID-19. In November 2021, the Omicron (BA.1) emerged in South Africa [11], another VOC that, according to GISAID in August 2022, had its sub-variants spreading to at least 193 countries [12,13]. The widespread and simultaneous circulation of both VOCs Omicron and Delta resulted in recombinants known as “Deltacron.” In France (February 2022), genomic analysis of SARS-CoV-2 samples pointed out a novel Deltacron lineage. This lineage presented two recombination breakpoints, one at the beginning of the spike region and another at the beginning of ORF3a (11). The genomic segment within these limits displayed Omicron signature mutations; however, the rest of the genome presented Delta signature mutations. This recombinant variant was then designated XD (12), mainly found in Denmark and the Netherlands (10,12,13). Besides XD, two other Omicron and Delta hybrids, XF and XS, circulated in the United Kingdom and the USA, respectively. Both recombinants presented a minor Delta portion at the 5′ end of an Omicron genomic backbone, although with distinct breakpoint locations (14). According to the Cov-Lineages [14], there are less than 40 sequences to each XD, XF, and XS recombinants.

Most recombinants available in the GISAD database were recovered from European countries and the USA. To fill the gap of studies in other regions, we investigated four putative Brazilian recombinants recovered from South and Southeast Brazil. We analysed the mutation profile and identified the recombination breakpoints, and we made a phylogenetic reconstruction to trace the origin of the novel recombinant lineages. This confirmed that the Deltacron recombinant in the country had evolved *de novo*, and that it can be considered a case of incipient parallel evolution. In other words, this recombinant variant acquired similar characteristics as other Deltacron variants (i.e., it shows a high level of sequence similarity), and the new variant AYBA-RS has evolved this independently from other circulating variants. Our results highlighted the importance of genomic surveillance for monitoring the viral evolution caused by coinfections with different SARS-CoV-2 lineages and for identifying putative recombinants. This is specifically pressing during periods of high viral circulation, and in countries with multiple variants, as well as in regions that are a hub for international air travel.

## Material and Methods

### Bioethics, sample collection and processing

The clinical samples were retrieved from three different institutions performing COVID-19 diagnosis and SARS-CoV-2 genomic surveillance in Rio Grande do Sul, Brazil: Centro Estadual de Vigilância em Saúde-CEVS (The State Centre for Health Surveillance from State Department of Health), Genetics and Molecular Biology Laboratory from Hospital Moinhos de Vento and Laboratório de Bioinformática Aplicada a Microbiologia Clínica from Federal University of Santa Maria (UFSM). In all cases, the SARS-CoV-2 infection was firstly detected by real time RT-PCR and the samples were submitted to genomic sequencing routine in each institution.

### Whole genome sequencing, assembly and quality control

We observed a *S* gene dropout (i.e. gene not detected) in the of the sample SC2-9898 on May 2022, and then selected this sample for genome sequencing with the SARS-CoV-2 FLEX NGS panel (PARAGON GENOMICS) on Illumina MiSeq platform. The library preparation was conducted according to the manufacturer’s protocol, and the sequencing was done using the MiSeq Reagent Micro Kit v2 (ILLUMINA). The FASTq files were obtained using the Local Run Manager Generate FASTQ Analysis Module v3.0 (ILLUMINA) and submitted to the SOPHiA DDM v.5 platform, which were analysed using the CleanPlex SARS hCoV2 pipeline for sequence alignment. Finally, the sequence was deposited in the GISAID database with the entry EPI_ISL_14381991.

For EPI_ISL_12110384 and EPI_ISL_14284846 sequences, the whole genome sequencing was performed using the Illumina COVIDSeq protocol (Illumina Inc, USA) on Illumina sequencing platform. The pipeline ViralFlow was used to perform genome assembly, variant calling, and consensus generation [15].

To evaluate the quality and determine the lineage of the genome sequences, we analysed them on Nextclade Web version 2.3.1 [16].

### Identification of lineage counterparts

To identify other genomes from the new Brazilian recombinant, we performed blast searches using the sequence Brazil/RS-FIOCRUZ-8390/2022 (EPI_ISL_12110384) on the “Unassigned” dataset from GISAID (assessed on 25 July 2022). After this, we visually inspected the mutation pattern of the top hits on the Nextclade Web, using the putative Brazilian recombinant sequences as reference.

### Parental lineages determination

Genome sequences were first aligned using the Nextalign version 1.11.0 with default parameters and sequence MN908947 as reference.

We evaluated the recombinant genomes using Sc2rf [17]. Subsequently, we manually segmented the genome of the oldest sequences of each recombinant lineage according to the Delta and Omicron portions using Aliview version 1.27 [18]. Then, we assessed the Pangolin lineage of each of the 5′ delta and the 3′ omicron segments in the Nextclade Web and Pangolin COVID-19 Lineage Assigner version 4.1.1 [19].

We also performed a blastn search (BLAST version 2.10.1+, [20,21]) of each of the Delta and Omicron segments on the sequences of Nextstrain’s global analysis - GISAID data [22], assessed on 23 June 2022). Next, we checked the lineage of the oldest top-hit strain in the GISAID metadata.

We built lineage-specific databases (GISAID sequences) considering the Pango lineages determined in the previous analyses. We again utilised each segment as a query to find the top 20 best hits of the reference databases (Delta or Omicron). Once we identified the best parental candidates, we downloaded their sequences from GISAID with the following filters: “low coverage excluded,”“collection date complete,” and “complete sequence.” Finally, we utilised these top hit sequences to compute each lineage’s frequency of mutations using a Python script (Pandas library version 1.4.2).

### Network analyses

To investigate the evolutionary history of the recombinants, we constructed a set that comprehended the Brazilian recombinant, XD, and XS lineages and their respective putative parental sequences. However, we only added the XF sequences to the dataset since we did not identify recombination in this lineage with the Sc2rf analysis (i.e., there were no candidate parental sequences to be included).

Subsequently, we aligned the sequences using Nextalign and submitted the dataset to a network analysis on Splitstree version 4.18.2 [23]. For this purpose, we used the NeighborNet method and drew the network using the RootedEqualAngle method using the Wuhan/WH01/2019 (EPI_ISL_406798) sequence as the root.

We also carried out a network analysis using the library pegas 1.1 from R version 4.1.3 [24]. We randomly sampled five sequences per lineage (AY.101, AY.4, B.1.617.2, BA.1, BA.1.1, XD, XS) from the original aligned dataset to improve the resolution of the network.

As before, we included all four sequences of the Brazilian recombinant in the sampled dataset. Finally, we determined the haplotypes using the function haplotype and carried out the network modelling using the haploNet method (default parameters).

### Recombination analyses

For recombination detection, we carried out two additional analyses. Firstly, we utilised the sampled dataset in the software RDP4 version 4.101 [25], using a “full exploratory recombination scan” (all methods with default parameters). Secondly, we performed an analysis with the HybridCheck R library version 1.0.1 [26]. For this analysis, we considered each segment’s oldest top hit to be the parental sequence.

Concerning the XF lineage, we used South Africa/NICD-N28358/2022 (Omicron) and South Africa/NHLS-UCT-GS-AF27/2021 (Delta) as the parental sequences, as described in Wang et al. (2022) [27].

Regarding the recombinant sequences described in this study, we annotated the genome mutations using the Coronapp [28]. We then drew the genome maps using the Python libraries, Seaborn and DNA features viewer 3.1.1 [29].

### Phylogenetic analyses

To investigate the phylogenetic history of the Brazilian recombinant segments, we concatenated the four identified sequences with their respective parental sequences (top hits of lineages AY. 101 and BA.1.1). We then split the aligned sequences into two segments, considering the recombination breakpoint inferred in the HybridCheck analysis: the 5′ portion encompassed positions 1-21769 and the 3′ portion, positions 21770-29903.

Next, we built a phylogenetic tree using IQ-Tree version 1.6.12 [30] with an automatically detected substitution model (option -m MFP) and 1,000 ultrafast bootstrapping replicates. Next, we conducted a timetree inference and a “mugration” model using discrete PANGO lineages with Treetime version 0.8.6 [31]. Subsequently, we drew a chronogram tree using a script written in R ggtree library version 3.2.1 [32], colouring the branches according to the PANGO lineages.

### Estimating the age of introgression

To infer an approximate date of the recombination event, we extracted the SNPs of the Brazilian recombinant with snp-sites version 2.5.1 [33], only outputting columns containing ACGT (option -c). We then calculated the coalescence time based on the formula described in Ward & Oosterhout (2015) [26], considering a mutation rate of 1.83 × 10-6 substitution per site per day [34] and genome size of 29,903 nucleotides, based on the reference genome (Wuhan/2019).

To evaluate the context of the cocirculating lineages in Brazil, we plotted a kernel density of absolute frequencies of Brazilian sequences collected between June 2021 and June 2022 (assessed in GISAID on 29 July 2022). We generated the density plots considering the Brazilian regions with a script written in Python (Seaborn library), kdeplot method with a smoothing parameter equal to 2 (bw_adjust=2). We assessed the association between the Brazilian regions (South and non-South) and Pango lineages (AY. 101, BA.1.1, and other lineages) using the Chi-square test (SciPy version 1.8.1). P-values <0.05 were considered statistically significant.

## Results

### Sampling, data acquisition and genome assembly

The SARS-CoV-2 recombinant samples were independently identified and processed by each institution according to their routine sequencing testing. The clinical data available and the assembly metrics for the three sequenced genomes are summarised in Supplementary Table S1.

### Identification of the Brazilian Deltacron, AYBA-RS

Preliminary analyses assigned the genome sequence from Cruz Alta (Brazil/RS-FIOCRUZ-8390/2022) to the recombinant lineage XS. However, the first 20 kb of the genome presented a distinct mutational pattern compared to an XS archetype (Supplementary Figure S1).

Through the genomic surveillance routine of the State Rio Grande do Sul, we identified two more sequences similar to the Cruz Alta sequence, one from Porto Alegre (Brazil/SC2-9898/2022) and another from Santa Maria (Brazil/RS-315-66266-219/2022) (Supplementary Table S1 and Supplementary Figure S2). Additionally, we searched the GISAID database and found a sequence from Rio de Janeiro (Brazil/RJ-NVBS19517GENOV829190059793/2022) that was very similar to the recombinants of South Brazil.

Once we identified our sequences as putative recombinants, we detected possible recombination signals in their genomes with Sc2rf. This analysis indicated that the 5′ region (positions 1-21845) came from a Delta lineage, and the 3′ region (positions 21846-29903) was from an Omicron lineage (Supplementary Figure S3). Further analyses indicated that the 5′ genomic region resembled mainly AY.101 and that from the 3′ region, BA.1 or BA.1.1 (Supplementary Table S2). Next, we built lineage-specific sequence databases and searched them for the most similar sequences to each segment (5′ Delta and 3′ Omicron). We considered the oldest top-hit for each segment to be the parental sequences, and we compared their mutational signatures to those of the Brazilian recombinant sequences (Figure 1A). In this analysis, all Brazilian recombinant sequences present similar patterns: their 5′ segment matched AY. 101, and their 3′ region, the BA.1.1 lineage (Figure 1 and Supplementary Figure S4). The substitution C10604T was found exclusively in all four sequences of the Brazilian recombinant (Supplementary Figure S4). Since the recombinant found in this study does meet the requirements of the Pango nomenclature [35], we named it as AYBA-RS, considering its parental lineages (<u>AY</u>.101 and <u>BA</u>.1.1) and the location of origin (<u>RS</u>, Brazil).

**Figure 1.**
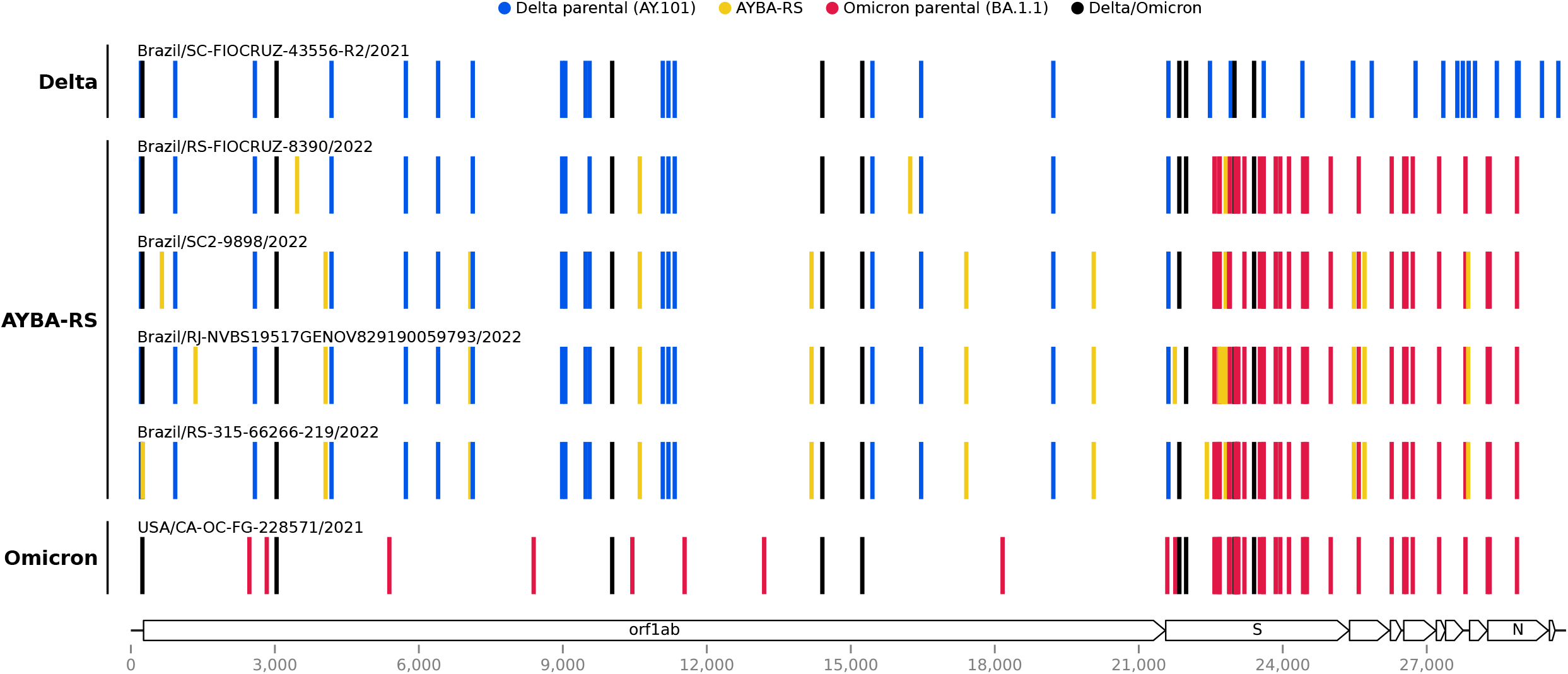
Mutation profiles of the Brazilian Deltacron and the Delta and Omicron variants. The Brazilian Deltacron presents a similar mutational pattern among the four Brazilian samples, where the first ~22 kb (5′ segment) resembles the Delta variant, and the last ~8 kb (3′ segment) the Omicron variant. Vertical colored lines represent nucleotide substitutions, with Wuhan/2019 as the reference. The legend depicts characteristic mutations of each parental sequence and of the recombinants. The SARS-CoV-2 genome map and their respective coordinates are shown at the bottom.

### Comparison between AYBA-RS and the other Deltacrons

We compared the AYBA-RS sequences to those from other Deltacrons described in Cov-Lineages [14], namely XD, XF, and XS. Identification of the recombination blocks using HybridCheck [26] supported the above result with Sc2rf, revealing a breakpoint at the beginning of the gene *S* (Figure 2, position 21769). Furthermore, the HybridCheck analysis showed that the recombination pattern differed from those of XD, XF, and XS (Figure 2). The XD and the AYBA-RS were mainly composed of a Delta scaffold, while the XF and XS of an Omicron one. Analysis with the RDP4 software [25] confirmed that the AYBA-RS arose from a single recombination event, separated from those that led to the other Deltacrons (Supplementary Table S3). This analysis also indicated a breakpoint close to the gene *S* (position 22675) (Supplementary Table S3), diverging by less than 1 kb concerning the HybridCheck breakpoint. The RDP4 analysis revealed recombination events for the XD and XS sequences, but not for XF sequences.

**Figure 2.**
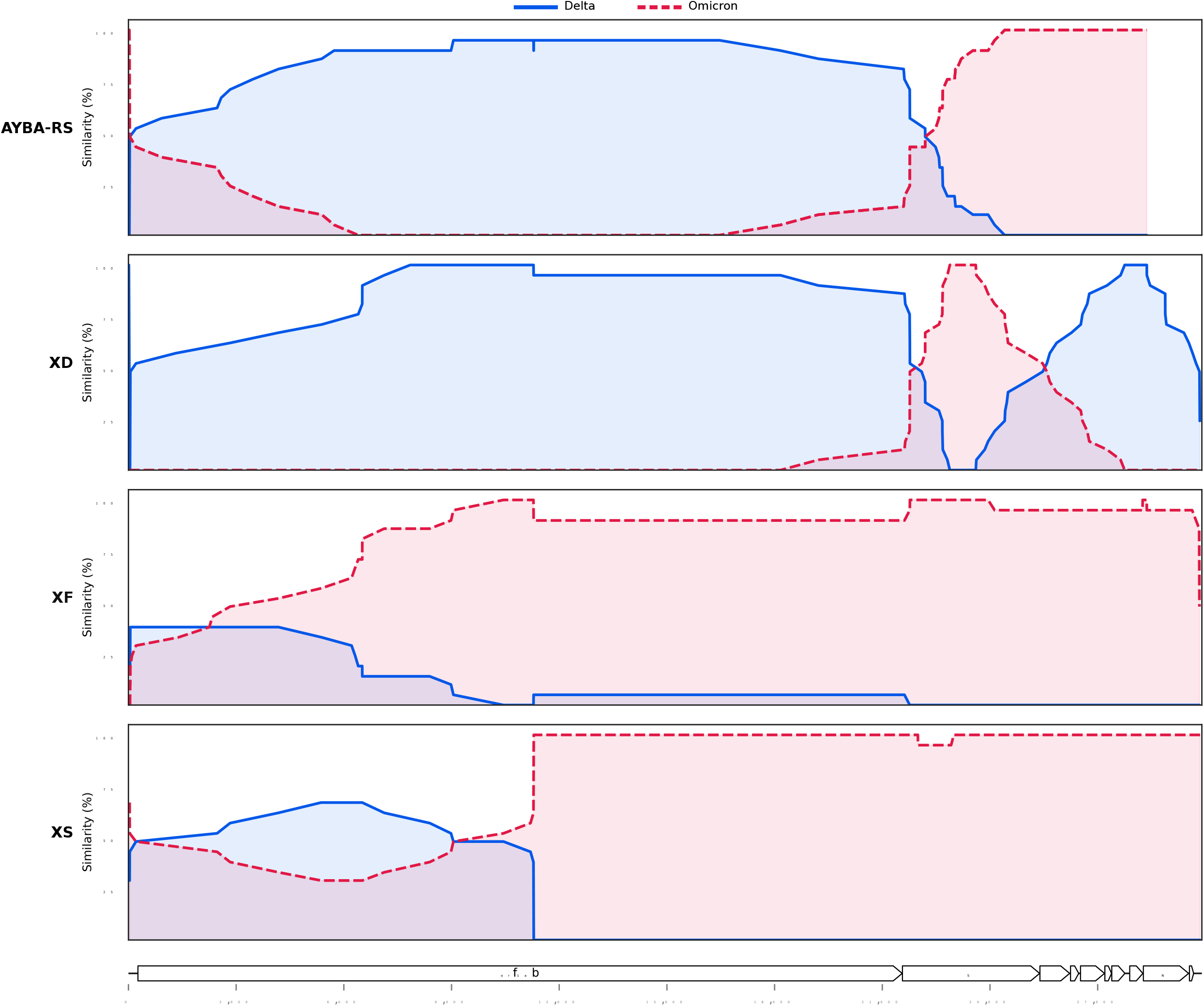
Comparison of the recombination patterns of the Brazilian Deltacron and the Deltacrons XD, XF, and XS. Similarity values between the recombinant and the Delta and Omicron parental sequences within a window length of 20 nucleotides along the genome. The legend describes the reference of each comparison. The recombination breakpoints, i.e., points where the lines cross over each other, diverge between the recombinants. Recombination plots generated with HybridCheck [26]. Sequences analysed in this plot: Brazil/RS-FIOCRUZ-8390/2022 (Brazilian Deltacron), Brazil/SC-FIOCRUZ-43556-R2/2021 (Delta) and USA/CA-OC-FG-228571/2021 (Omicron) as its parental sequences; France/HDF-IPP54794/2022 (XD), Sweden/37524448XXP/2021 (Delta) and Finland/P-1301/2022 (Omicron) as its parental sequences; England/PHEC-YYN8J41/2022 (XF), South Africa/NHLS-UCT-GS-AF27/2021 (Delta) and South Africa/NICD-N28358/2022 (Omicron) as its parental sequences; USA/CO-CDC-FG-248528/2022 (XS), Latvia/3410639/2021 (Delta) and USA/CA-CDC-FG-223742/2021 (Omicron) as its parental sequences. Similarity at the y-axis refers to percentage sequence similarity at polymorphic sites only. The SARS-CoV-2 genome map and their respective coordinates are shown at the bottom.

### Evolutionary history of recombinants of VOC Delta and VOC Omicron

Phylogenetic network reconstruction and haplotype network analysis were congruent since all four Brazilian recombinant sequences formed a distinct group from the other Deltacrons. Furthermore, both models showed that the different Deltacrons were distributed between the Delta and Omicron groups, having additional portions of each lineage (Figure 3A and Figure 3B).

**Figure 3.**
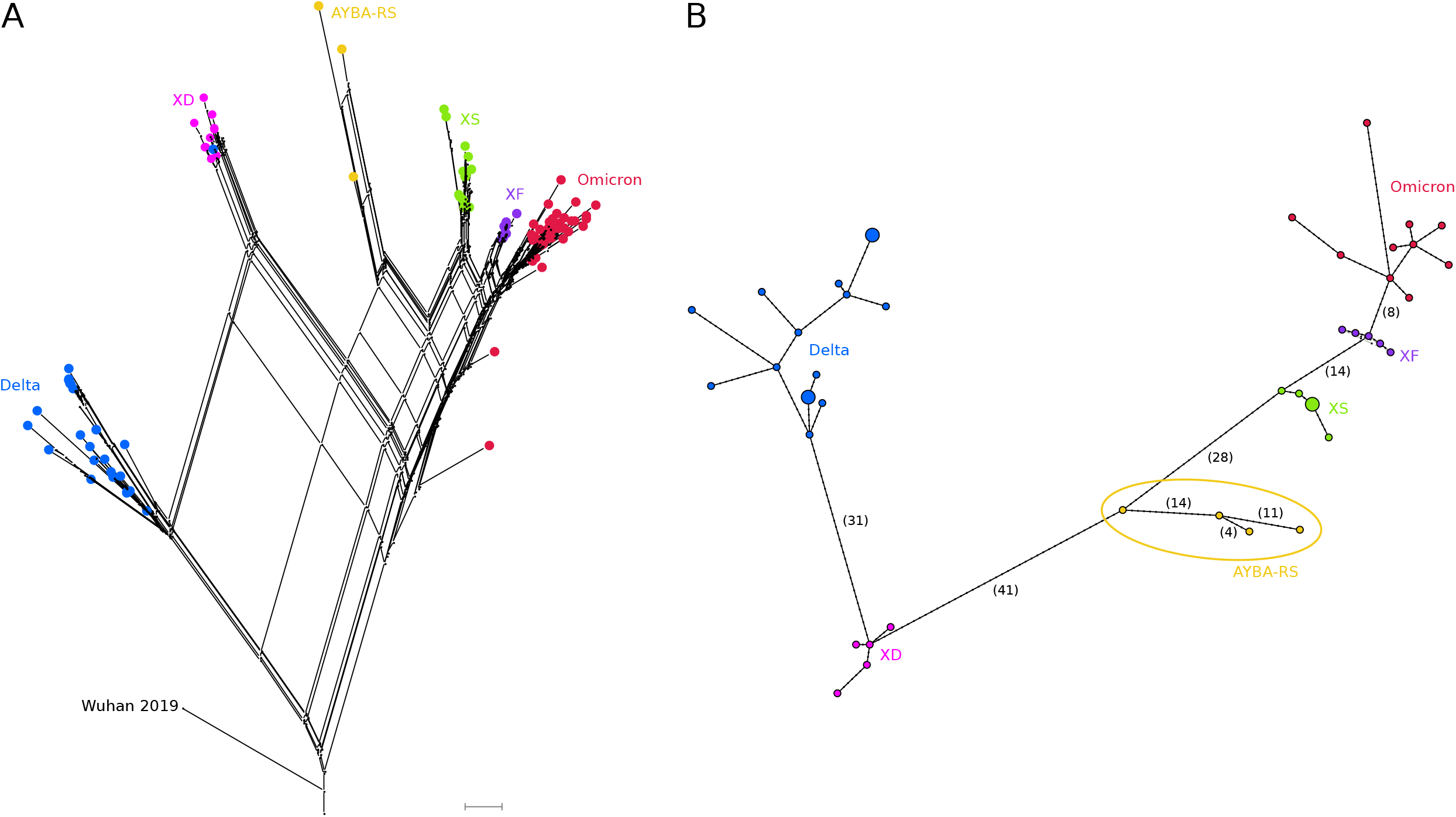
Evolutive history of Deltacron variants. **A** - Phylogenetic network of Deltacron sequences built with SplitsTree (NeighborNet method, RootedEqualAngle on Wuhan/WH01/2019). **B** - Haplotype network of Deltacron sequences made with pegas (haploNet method). The number of SNPs is in parentheses (Supplementary Table S5); the AYBA-RS is within an ellipse. In both analyses, Delta (AY.101, AY.4, B.1.617.2) and Omicron (BA.1, BA.1.1) sequences were used as references. Nodes are coloured according to the variant type.

Phylogenetic analyses for each of the 5′ (Delta) and 3′ (Omicron) blocks of the AYBA-RS, using the best matches to the AY. 101 and BA.1.1 lineages, assigned the 5′ segments of the Brazilian Deltacron to the AY.101 clade. This clade was formed only by sequences from Brazil, notably from Santa Catarina (SC), a State from the South region that borders the Rio Grande do Sul (RS) (Supplementary Figure S5). On the other hand, the 3′ segments of the AYBA-RS formed a clade with BA.1.1 sequences from diverse geographical locations. However, in this tree, the AYBA-RS did not form a group with sufficient bootstrap support (Supplementary Figure S5).

Considering the number of SNPs between the AYBA-RS sequences (Figure 2B, Supplementary Table S4 and Supplementary Table S5), we estimated that the recombination event that gave origin to the recombinant may have occurred 180 days before the collection date of the first sample, i.e., December 2021. Inspection of the lineage density plots revealed an overlap of AY.101 and BA.1.1, mainly in December 2021, across the country’s regions (Figure 4). AY.101 and BA.1.1 presented higher relative frequency in the South region than in the rest of Brazil (Supplementary Table S6, Chi-square test: *X*^2^ = 10519.21, d.f.=1, p < 0.00001).

**Figure 4.**
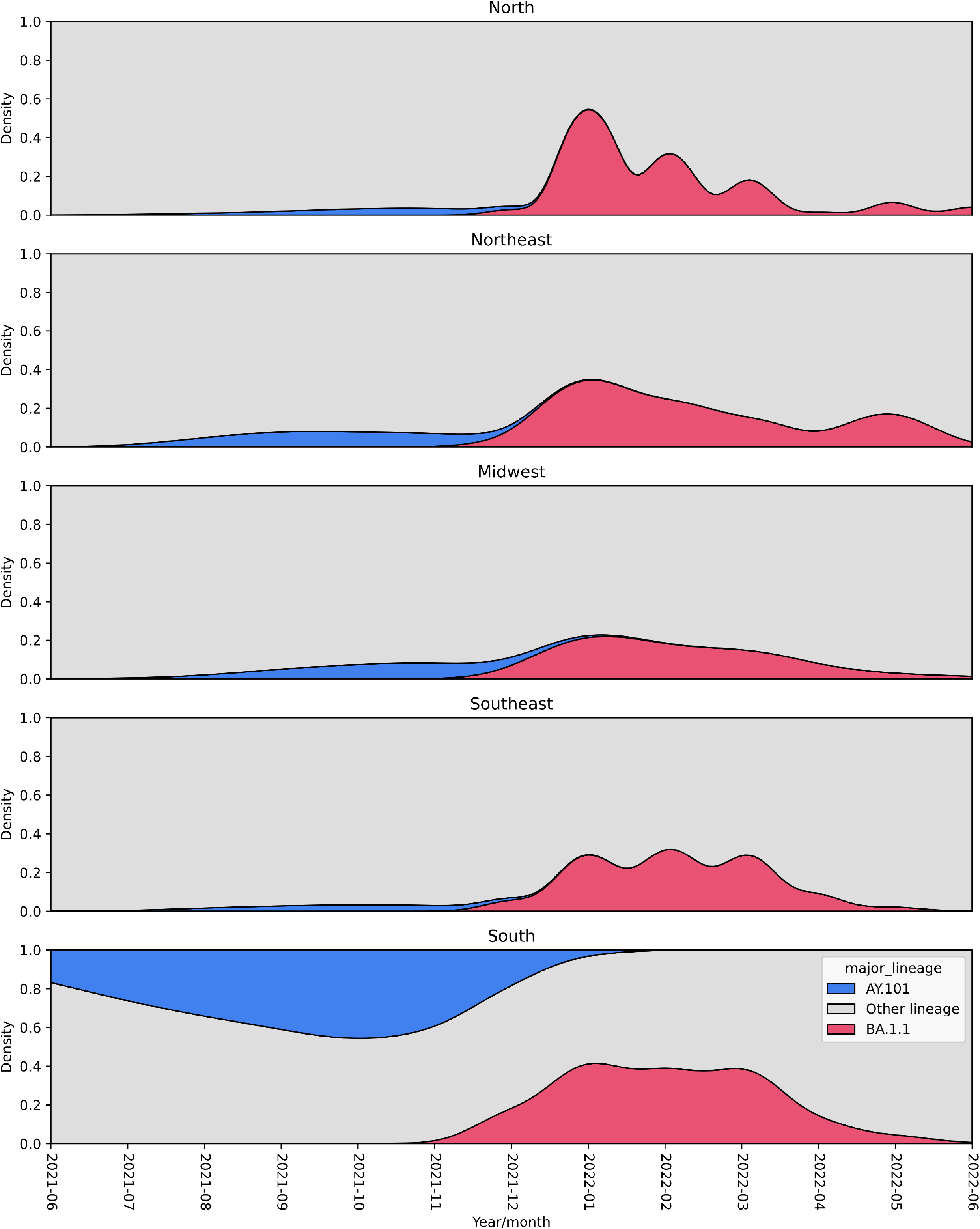
Density plot of the frequencies of AY.101 (Delta) and BA.1.1 (Omicron) lineages circulating in Brazil between June 2021 to June 2022. Both lineages co-circulated mainly in December 2021 in South Brazil. The colour of the lineages is shown in legend.

## Discussion

Here we describe the first Deltacron lineage identified in Brazil, AYBA-RS. Our analysis shows that this recombinant strain arose from a single recombination event between the AY. 101 and the BA.1.1 lineages in Southern Brazil. The genetic exchange between both variants most likely happened in December 2021, when the Omicron lineage started to take over Delta around the country [36]. Furthermore, we show that this recombinant differs from the previously described SARS-CoV-2 Deltacrons XD, XF, and XS, supporting a new recombination event and evidence of incipient parallel evolution. We employed a robust approach to identify and describe the recombinant SARS-CoV-2 lineages, combining methods involving phylogenetic and population genetic techniques incorporated in HybridCheck [26], RDP4 [25], SplitsTree [23] and other approaches. This combined approach enabled us to determine the parental lineages, identify the recombination breakpoint around the 22 kb position near the Spike gene (*S*), and estimate the age of the new recombinant lineage.

Based on the phylogenetic reconstruction we were able to ascertain that the sequences found in Rio Grande do Sul and Rio de Janeiro states coalesced and have a single origin. Since genomic deposits in GISAID are recent, the lack of more sequences in the database suggests that the variant had a minor epidemiological impact. Alternatively, a lack of genomic surveillance may also have contributed. Indeed, it is essential to consider undersampling bias since the sequencing effort from Brazil is still lower than those from Europe and the USA [37]. Therefore, it is not surprising that most of the recombinants described are from Europe and the USA, although we cannot rule out that this could also reflect a genuine pattern, given that these regions experience considerable international air travel.

Co-infections with different variants of SARS-CoV-2 are necessary to trigger recombination events. Spatiotemporal variation in selection pressures can maintain a balanced polymorphism and maintain multiple variants. In addition, multiple variants can also be maintained in substructured environments, as well as by a time-lag in coevolution (see Introduction). International travel can mediate gene flow and bring these distinct variants into contact, thereby facilitating inter-variant recombination and genetic introgression [4]. In fact, along the pandemic course, there have been reports of patients having Omicron and Delta co-infections [38–40]. Such events provide an opportunity for the emergence of new lineages with distinct phenotypes [3,41]. These phenotypes can occupy different peaks in the fitness landscape separated by fitness valleys. Such valleys can be a consequence of epistasis, which is a phenomenon wherein nucleotide substitutions influence each other’s impact on fitness. This results in a fitness landscape with many small and large peaks, ridges, and valleys. In such a rugged landscape, populations evolve slowly because they can become stuck once they have reached a local optimum, i.e., the highest fitness peak in the nearby landscape [42]. In that case, several mutations are required to climb the next even higher peak [43]. Recombination events could help the virus to bridge such valleys because recombination offers three theoretical advantages over mutations (see Introduction). Given the large amount of nucleotide divergence that has evolved in multiple extant lineages, we argue that it is likely that recombinant evolution will play an increasingly important role in SARS-CoV-2 evolution and the COVID-19 pandemic. The potential for recombination to evolve better adapted SARS-CoV-2 variants is increased by international travel that can bring allopatric lineages and variants from different continents together.

An analysis proposed by Turakhia and co-workers [44] suggested that approximately 2.7% of sequenced SARS-CoV-2 genomes have detectable recombinant ancestry. However, the authors also highlight that hybrid strains of genetically similar viral lineages are challenging to detect, and that the overall recombination frequency could be underestimated [44]. The current study corroborates this assertion, showing that distinct recombinant lineages can be challenging to differentiate, and that advanced evolutionary genomic analyses are required to identify and trace back the origin of recombinant lineages.

The genomic bulletin from June 2022, which included only 83 samples collected in the Rio Grande do Sul, revealed that even with the predominance of the Omicron lineage, Delta (AY.99.2) and Gamma (P.2) lineages are still circulating. Taking into account the relaxation of prevention measures, the non-adherence to the vaccine booster dose, and the simultaneous circulation of multiple lineages in the same region, we might be creating a perfect storm for the emergence of new SARS-CoV-2 variants of concern. Our study supports the assertion that SARS-CoV-2 genetic introgression events might be more common than expected initially. This has implications for disease control measures, and it emphasises the need for more intensive genomic and epidemiological surveillance across the world.

## Acknowledgments

We thank M.Sc. Luana Giongo Pedrotti for statistical support, and Edmund Willis (MD) for commenting on an earlier draft of the MS. We thank all the authors who have shared genome data on GISAID utilised in this study: all genome sequences and associated metadata are published in GISAID’s EpiCoV database (EPI_SET_220829tz). To view the contributors of each sequence with details such as accession number, Virus name, Collection date, Originating Lab and Submitting Lab, and the list of Authors, visit 10.55876/gis8.220829tz. CVO is funded by the University of East Anglia (UEA) and Earth and Life Systems Alliance (ELSA), Norwich Research Park, Norwich, UK.

## Supplementary Figures

**Supplementary Figure S1. Mutational patterns of the Brazilian and the XS recombinants (Nextclade’s output).** The Brazilian recombinant was initially identified as an XS lineage, although the portion between the positions 1 and 20000 diverged from the XS archetype.

**Supplementary Figure S2. Origin of the Brazilian Deltacron samples.** Map of Brazil showing where the Brazilian Deltacron samples were collected (yellow points) with their respective collection dates. GISAID accession numbers are in parentheses.

**Supplementary Figure S3. Recombination detection in the genome sequence of Brazil/RS-FIOCRUZ-8390/2022 (Sc2rf output).** The analysis indicated that the 5′ region (positions 1-21845) came from Delta, and the 3′ region (positions 21846-29903) came from an Omicron.

**Supplementary Figure S4. Nucleotide assignatures of each lineage.** Heatmap showing the frequency of nucleotide mutations in each lineage. We used the blastn top-hits sequences for each parental lineage AY. 101 (n=20) and BA.1.1 (n=19). Mutation frequencies of AYBA-RS were computed from the 4 sequences found in Brazil. Asterisk - exclusive mutation of AYBA-RS.

**Supplementary Figure S5. Phylogenetic history of genomic segments of the Brazilian Deltacron.** A - The 5′ segment (coordinates 1-21769) of the Brazilian recombinant grouped to the Brazilian AY.101 genomes. B - The 3′ segment (coordinates 21770-29903) of the Brazilian recombinant grouped to BA.1.1 genomes from diverse geographical locations. Ultrafast bootstrap values of the main branches are close to the nodes.

## Supplementary Tables

Supplementary Table S1. Clinical data of the samples and characteristics of the genomes sequenced in this study.

Supplementary Table S2. Lineage identification of the segments of the earliest Deltacron sequences

Supplementary Table S3. Recombination analysis of Deltacron variants (RDP4 output).

Supplementary Table S4. Time of divergence between the AYBA-RS genome sequences.

Supplementary Table S5. SNPs of the AYBA-RS genome sequences.

Supplementary Table S6. Percentage of AY.101, BA.1.1, and other lineages between the South and the rest of Brazil.

