## Supplementary file for "Incipient parallel evolution of SARS-CoV-2 Deltacron variant in South Brazil"

3 - Centro de Desenvolvimento Científico e Tecnológico, Centro Estadual de Vigilância em Saúde, Secretaria Estadual da Saúde do Rio Grande do Sul (CDCT/CEVS/SES-RS), Porto Alegre, Rio Grande do Sul, Brazil

4 - Laboratório Central de Saúde Pública, Centro Estadual de Vigilância em Saúde, Secretaria Estadual da Saúde do Rio Grande do Sul (LACEN/CEVS/SES-RS), Porto Alegre, Rio Grande do Sul, Brazil

5 - Laboratory of Respiratory Viruses and Measles, Oswaldo Cruz Institute (IOC), Oswaldo Cruz Foundation (FIOCRUZ), Rio de Janeiro, RJ, Brazil

6 - Departamento de Entomologia e Núcleo de Bioinformática, Instituto Aggeu Magalhães, Fundação Oswaldo Cruz Pernambuco (FIOCRUZ-PE), Recife, Pernambuco, Brazil

7 - Universidade Federal de Santa Maria, Santa Maria, RS, Brazil

8 - School of Environmental Sciences, University of East Anglia, Norwich Research Park, Norwich, UK

***Corresponding authors:** Eliana Wendland and Cock van Oosterhout

### Supplementary Figures

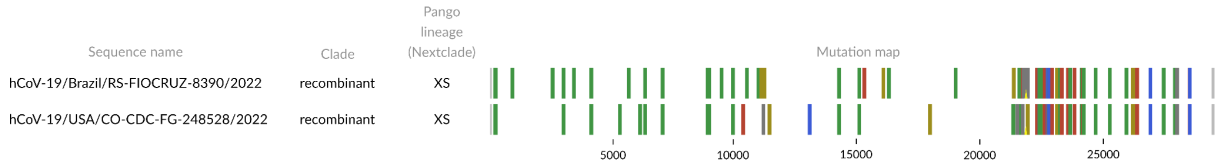


**Supplementary Figure S1. Mutational patterns of the Brazilian and the XS recombinants (Nextclade's output).** The Brazilian recombinant was initially identified as an XS lineage, although the portion between the positions 1 and 20000 diverged from the XS archetype.

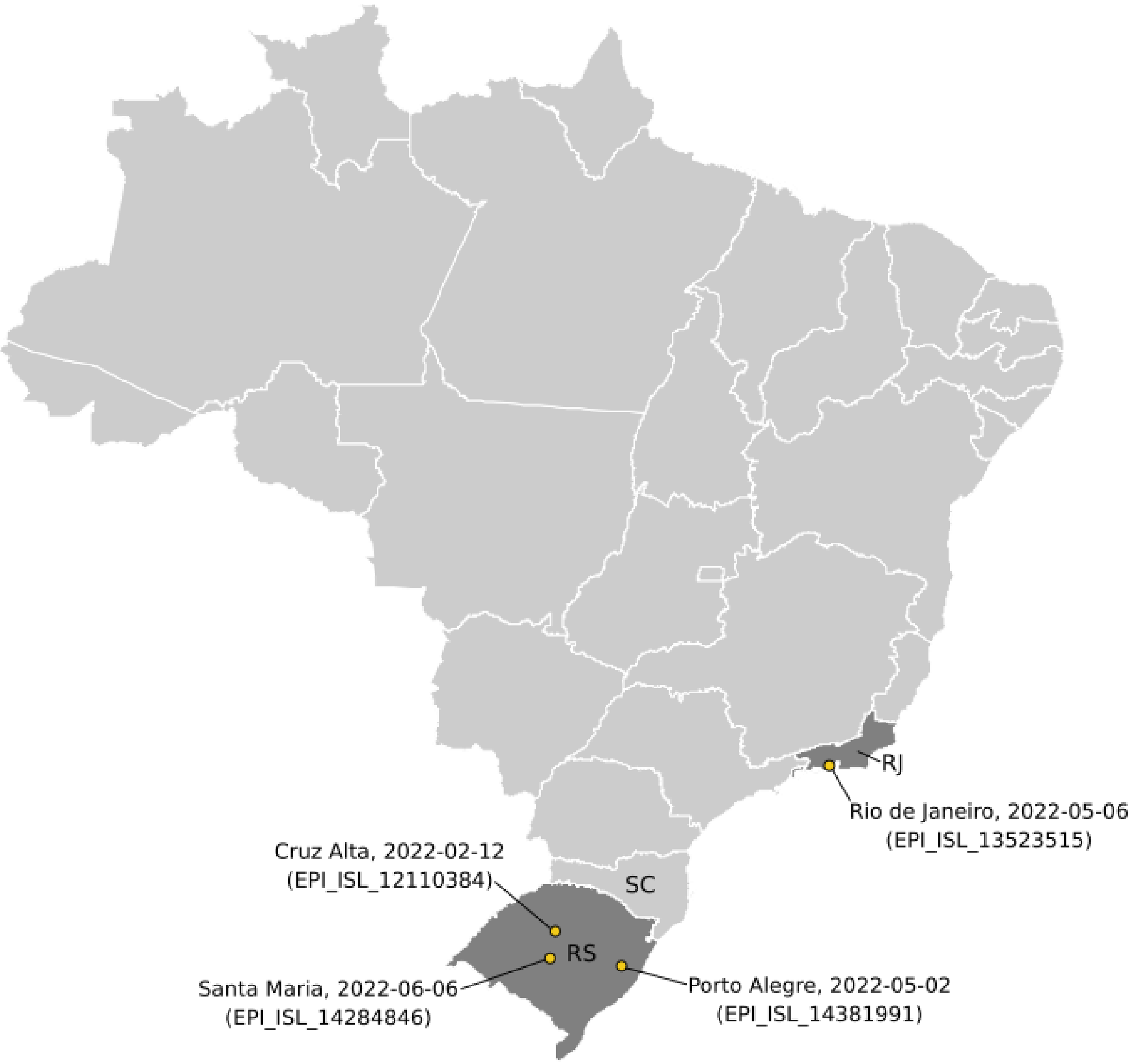


**Supplementary Figure S2. Origin of the Brazilian Deltacron samples.** Map of Brazil showing where the Brazilian Deltacron samples were collected (yellow points) with their respective collection dates. GISAID accession numbers are in parentheses.

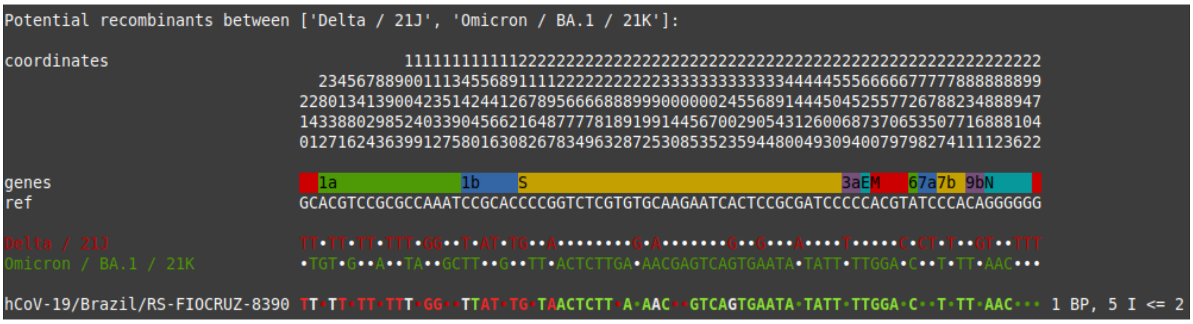


**Supplementary Figure S3. Recombination detection in the genome sequence of Brazil/RS-FIOCRUZ-8390/2022 (Sc2rf output).** The analysis indicated that the 5' region (positions 1-21845) came from Delta, and the 3' region (positions 21846-29903) came from an Omicron.

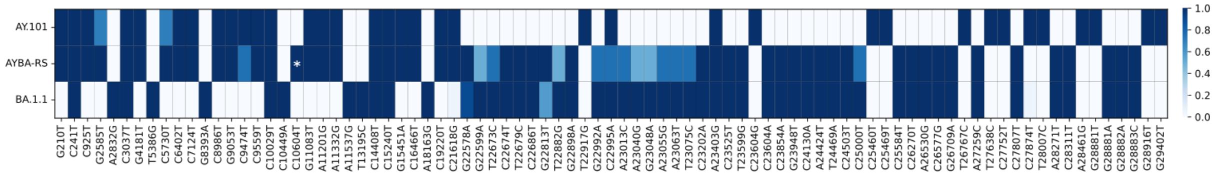


**Supplementary Figure S4. Nucleotide assignatures of each lineage.** Heatmap showing the frequency of nucleotide mutations in each lineage. We utilized the blastn top-hits sequences for each parental lineage AY.101 (n=20) and BA.1.1 (n=19). Mutation frequencies of AYBA-RS were computed from the 4 sequences found in Brazil. Asterisk - exclusive mutation of AYBA-RS.

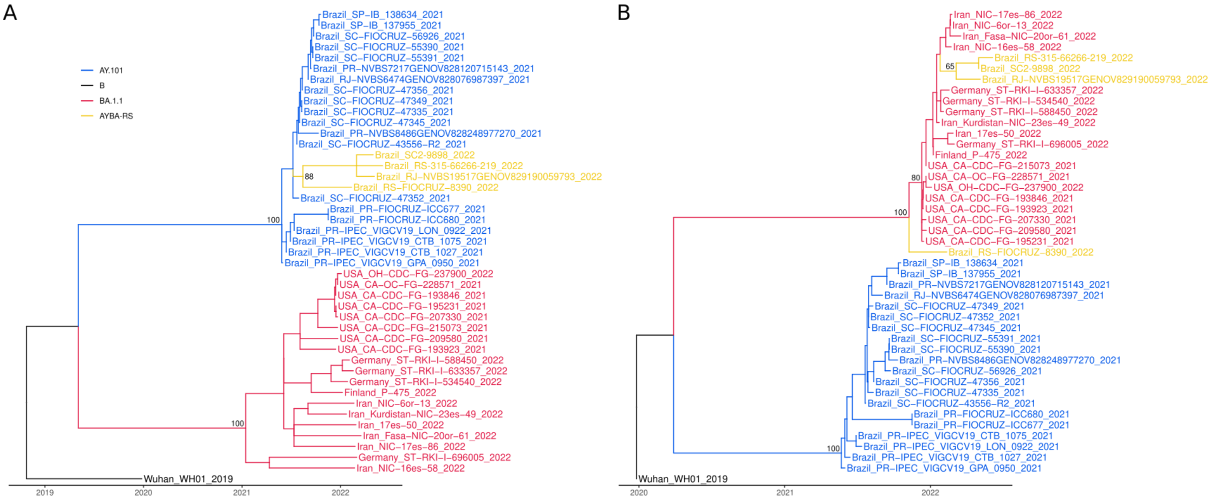


**Supplementary Figure S5. Phylogenetic history of genomic segments of the Brazilian Deltacron.** A - The 5' segment (coordinates 1-21769) of the Brazilian recombinant grouped to the Brazilian AY.101 genomes. B - The 3' segment (coordinates 21770-29903) of the Brazilian recombinant grouped to BA.1.1 genomes from diverse geographical locations. Ultrafast bootstrap values of the main branches are close to the nodes.

### Supplementary Tables

**Supplementary Table S1. Clinical data of the samples and characteristics of the genomes sequenced in this study.**

| | Brazil/RS-315-66266-219/2022 | Brazil/SC2-9898/2022 | Brazil/RS-FIOCRUZ-8390/2022 |
|---|---|---|---|
| **Accession ID** | EPI_ISL_14284846 | EPI_ISL_14381991 | EPI_ISL_13523515 |
| **Sample date** | 06/06/2022 | 02/05/2022 | 12/02/2022 |
| **Location** | Santa Maria | Porto Alegre | Cruz Alta |
| **Mapped reads** | 514,768 | 602,459 | 3,039,386 |
| **Coverage breadth** | >30x 98.79% | >30x 98.70% | >30x 99% |
| **Coverage depth** | 798x | 532x | 10000x |
| **Library construction method** | COVIDSeq Assay Illumina | PARAGON CleanPlex SARS-CoV-2 FLEX Panel | COVIDSeq Assay Illumina |
| **Sequencing technology** | Illumina iSeq 100 | Illumina MiSeq | Illumina MiSeq |
| **Assembly method** | ViralFlow | SOPHiA DDM v.5 | ViralFlow |

**Supplementary Table S2. Lineage identification of the segments of the earliest Deltacron sequences**

| Sample (Lineage) | Segment start | Segment end | Nextclade* | Pangolin# | Lineage of the top-hit (strain name)$ |
|---|---|---|---|---|---|
| Brazil/RS-FIOCRUZ-8390/2022 (AYBA-RS) | 37 | 21845 | AY.101 | AY.101 | AY.100 (Guatemala/INC-LNS-127/2021) |
| Brazil/RS-FIOCRUZ-8390/2022 (AYBA-RS) | 21846 | 29857 | BA.1 | - | BA.1.1 (Taiwan/TSGH-52/2021) |
| France/HDF-IPP54794/2022 (XD) | 55 | 21845 | XD | XD | BA.1 (Brazil/BA-FIOCRUZ-PVM99977/2022) |
| France/HDF-IPP54794/2022 (XD) | 21846 | 25469 | XD | - | AY.4 (Belgium/ULG-17464/2021) |
| USA/CO-CDC-FG-248528/2022 (XS) | 38 | 10029 | XS | - | B.1.617.2 (Pakistan/UHSPK3-61/2021) |
| USA/CO-CDC-FG-248528/2022 (XS) | 10030 | 29792 | XS | - | BA.1.1 (Paraguay/454211/2022) |

* - Lineage identification in the Nexclade Web

\# - Lineage identification in the Pangolin COVID-19 Lineage Assigner

$ - Blast on Nextstrain's global analysis - GISAID data, lineage taken from the GISAID metadata

**Supplementary Table S3. Recombination analysis of Deltacron variants (RDP4 output).**

| Recombinant | Breakpoint Start | Breakpoint End | Minor parental lineages | Major parental lineages | RDP (p-value) | GENECONV (p-value) | Maxchi (p-value) | Chimaera (p-value) | SiSscan (p-value) |
|---|---|---|---|---|---|---|---|---|---|
| AYBA-RS | 22675 | 29392 | BA.1.1, BA.1, XF, XS | AY.4, AY.101, B.1.617.2 | 8.21E-06 | 3.89E-05 | 6.05E-10 | 1.20E-09 | 1.33E-08 |
| XD | 21804 | 25526 | BA.1.1, BA.1, XF, XS | AY.4, AY.101, B.1.617.2 | 1.76E-08 | 5.11E-09 | 1.33E-09 | 6.12E-10 | 2.45E-07 |
| XS | 29652 | 9751 | AY.4, AY.101, B.1.617.2, XD | BA.1.1, BA.1 | 0.012 | 2.04E-04 | 0.001 | 0.002 | NS |

NS - non-significant

**Supplementary Table S4. Time of divergence between the AYBA-RS genome sequences.**

| Sequences | SNP divergence | Mean time (in days) | Min time (5% CI) | Max time (95% CI) |
|---|---|---|---|---|
| CA – RJ | 19 | 180 | 120 | 255 |
| SM – CA | 16 | 152 | 100 | 222 |
| PA – CA | 14 | 134 | 84 | 200 |
| SM – RJ | 13 | 125 | 78 | 190 |
| PA – RJ | 11 | 107 | 63 | 168 |
| PA – SM | 4 | 42 | 18 | 83 |

CA - Brazil/RS-FIOCRUZ-8390/2022 (collection date, 2022-02-11)

PA - Brazil/SC2-9898/2022 (collection date, 2022-05-02)

RJ - Brazil/RJ-NVBS19517GENOV829190059793/2022 (collection date, 2022-05-06)

SM - Brazil/RS-315-66266-219/2022 (collection date, 2022-06-06)

### Supplementary Table S5. SNPs of the AYBA-RS genome sequences.

| Position | CA | PA | RJ | SM |
|---|---|---|---|---|
| 245 | C | C | C | T |
| 647 | A | G | A | A |
| 1348 | C | C | T | C |
| 3464 | T | C | C | C |
| 4057 | T | C | C | C |
| 7075 | T | C | C | C |
| 7081 | C | T | T | T |
| 14183 | C | T | T | T |
| 16238 | G | C | C | C |
| 17407 | T | C | C | C |
| 20062 | G | T | T | T |
| 21752 | T | T | C | T |
| 21846 | T | T | C | T |
| 22419 | C | C | C | T |
| 22599 | G | A | G | A |
| 22673 | C | C | T | C |
| 22688 | A | A | G | A |
| 22775 | G | G | A | G |
| 22786 | A | A | C | A |
| 22792 | C | C | T | C |
| 22882 | T | G | T | G |
| 25000 | T | T | T | C |
| 25482 | C | A | A | A |
| 25704 | T | C | C | C |
| 27864 | C | T | T | T |

CA - Brazil/RS-FIOCRUZ-8390/2022 (collection date, 2022-02-11)

PA - Brazil/SC2-9898/2022 (collection date, 2022-05-02)

RJ - Brazil/RJ-NVBS19517GENOV829190059793/2022 (collection date, 2022-05-06)

SM - Brazil/RS-315-66266-219/2022 (collection date, 2022-06-06)

**Supplementary Table S6. Percentage of AY.101, BA.1.1, and other lineages between the South and the rest of Brazil.**

| Lineage | South Brazil | Non-South Brazil |
|---|---|---|
| AY.101 | 15.68%<br>(2580) | 1.44%<br>(1590) |
| BA.1.1 | 16.08%<br>(2646) | 8.45%<br>(9344) |
| Other lineages | 68.24%<br>(11228) | 90.11%<br>(99645) |

Absolute frequencies are in parentheses.
